# RECRUITMENT OF GROUPER BROODSTOCK ON THE BASIS OF SINGLE LOCUS DNA MARKERS

**DOI:** 10.1101/067231

**Authors:** Kenneth F. Rodrigues, Ahmad Z. Tani, Syarul N. Baharum

## Abstract

Scientific breeding programs are founded on the screening and recruitment of genetically diverse broodstock, with the ultimate aim of developing heterogeneous breeding populations that host a collection of desirable traits. Single locus DNA markers can be applied to facilitate the process of selection as they are species specific, reliable, reproducible and easy to use. This study set forth to develop a library of single locus DNA markers for two commercially cultured species of groupers, *Epinephelus fuscoguttatus* and *E. corallicola*. DNA was isolated from one representative specimen of each species and utilized to construct shotgun genomic libraries. DNA sequences derived from the library were selected for the development of 42 and 41 single locus DNA markers for *E. fuscoguttatus* and *E. corallicola* respectively. The markers were then tested against randomly selected specimens obtained from the wild. Genotyping results revealed that the species specific primers demonstrated the ability to distinguish between individuals from the same species into distinct operational taxonomic units (OTUs) on the basis of their differential DNA profiles, thus establishing a basis for selection based on genetic heterogeneity. The findings of this study present a strong case for the application of single locus DNA markers as molecular tools for the selection of broodstock on the basis of genotyping.

## 1.0 INTRODUCTION

The economic success of the aquaculture industry is founded on the selection and recruitment of high quality broodstock from genetically diverse wild populations. However, the process of selection of wild specimens is confounded by phenotypical similarity and paucity of genetic information. Genomic molecular markers [1] have the potential to revolutionize the aquaculture industry when applied in conjunction with conventional breeding techniques [2]. This has formed the basis for approaches that recruit broodstock and seed on the basis of data obtained from Quantitative Trait Loci (QTLs) [3] and Marker Assisted Selection (MAS) [4]. These two molecular approaches can have a significant impact on the economics of a fish breeding operation as they offer an avenue for the elimination of genetically inferior germplasm prior to recruitment. Scientists have initiated the process of marker development for finfish by focusing on several commercially exploited species such as *Orecohromis niloticus* [5], *Ictalurus punctatus* [6], *Salmo salar* [7] and *Cyprinus carpio* [8]. They have relied on an assortment of molecular markers such as microsatellites [9], Single Nucleotide Polymorphisms (SNPs) [10], Amplified Fragment Length Polymorphisms [11] and Short Sequence Repeats [12] in order to detect genetic polymorphisms that can be exploited by breeders. The practical application of molecular markers to breeding has been discussed in detail [13] and it is interesting to note that highly informative markers such as microsatellites and SNPs require specialized laboratories and analytical techniques, thus putting them beyond the technological reach of commercial breeders. Single locus DNA markers [14] are less informative than microsatellites and SNPs, however they make up for this shortcoming by their simplicity, reproducibility and ability to distinguish individuals within a population. This study set forth to develop single locus DNA markers could be applied for the selection of broodstock purely on the basis of Boolean DNA profiling. The groupers *E. fuscoguttatus* and *E. corallicola* were selected as the target species as they are commercially exploited by the aquaculture industry.

## 2.0 MATERIAL AND METHODS

### 2.1 Sample collection and DNA isolation

The samples for this study were all collected from broodstock being maintained at the Borneo Marine Research Institute, Universiti Malaysia Sabah. DNA was extracted from Fin clips collected from *E. fuscoguttatus* (12) and *E. corallicola* (10) using the DNeasy extraction kit (Qiagen), the concentration of DNA was assessed using a micro-volume UV-Vis Spectrophotometer (GE Healthcare Life Sciences) and the final concentration was adjusted to 50 ng/μl using sterile nuclease free water. DNA samples were stored at −80° C.

### 2.2 Construction of genomic libraries

Partial small insert genomic libraries [15] were constructed using one DNA sample derived from each of the two species of groupers, *E. fuscoguttatus* and *E. corallicola*. Restriction digests were carried out using a combination of six restriction enzymes, *Eco*RI/*Hind*III, *Eco*RI/<u>*Bam*</u>HI, *Bam*HI/*Hind*III, *Kpn*I/*Sal*I, *Sac*I/*Kpn*I and *Sac*I/*Xba*I. (New England Biolabs), following which the digested DNA was ligated onto a pUC19 Vector and transformed into chemically competent *Escherichia coli* (TOP10). Recombinant clones from each library were selected randomly from Luria-Bertani plates containing Ampicillin (100 mg/l) and X-Gal (50 mg/l). Plasmid DNA were extracted and purified using GeneJET plasmid purification kit (Fermentas) and sequenced using BigDye Terminator 2.0 Cycle Sequencing Ready Reaction Kit (Life Technologies, USA) on an ABI Prism 3770 automated DNA sequencer (Applied Biosystems). Sequences were deposited at the NCBI GenBank and assigned accession numbers (Tables 1.0 and 3.0).

**Table 1.0:**
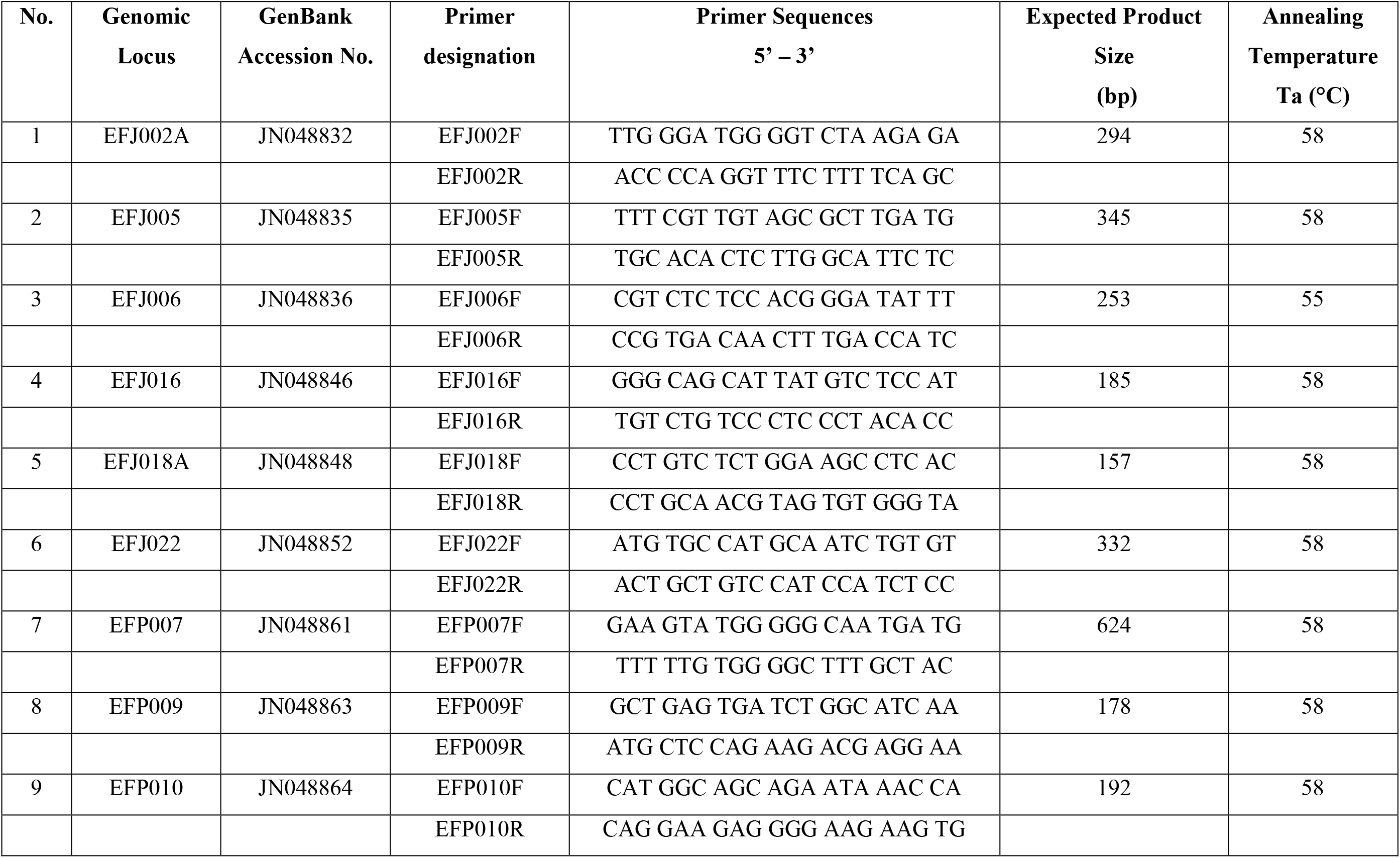
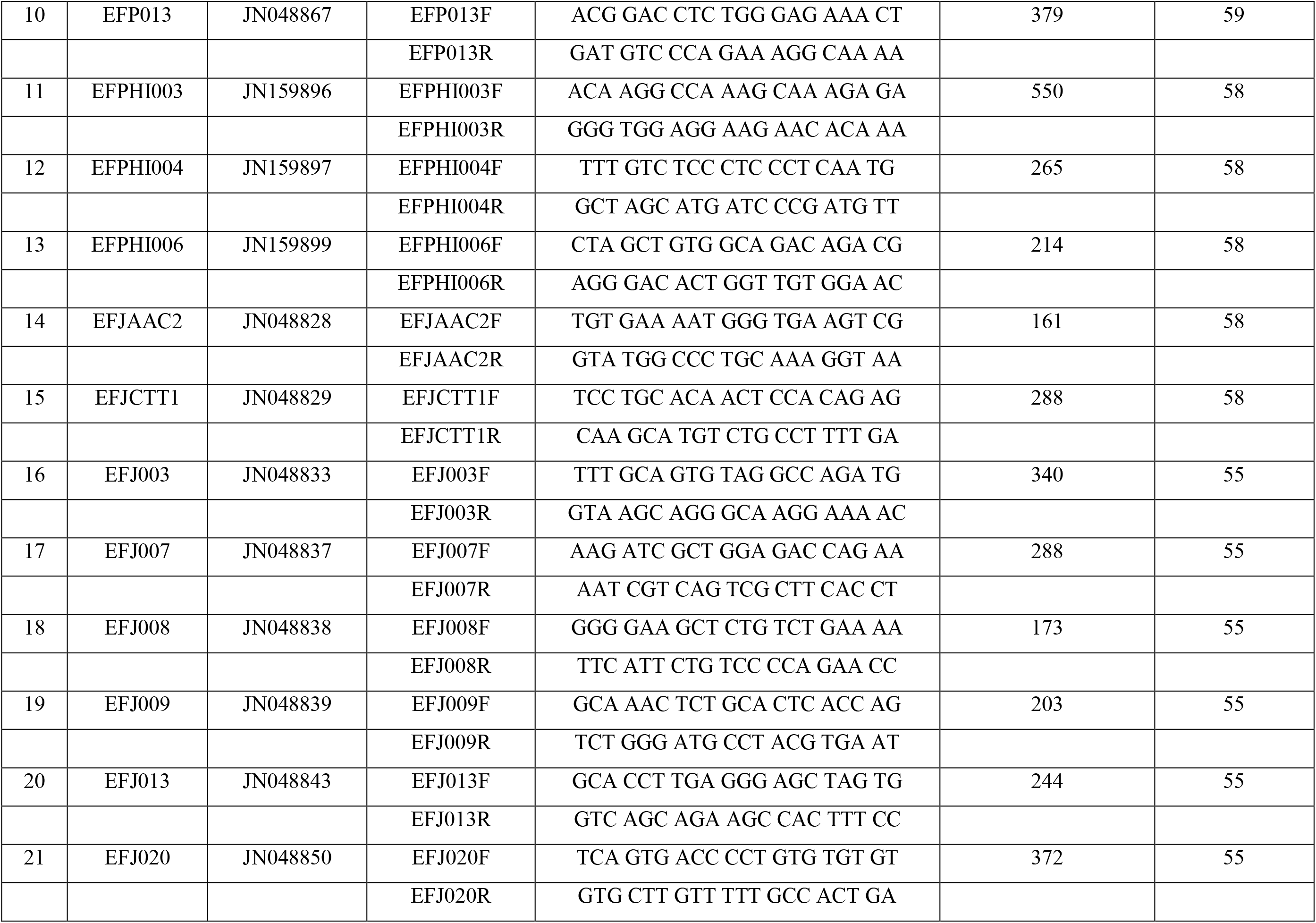
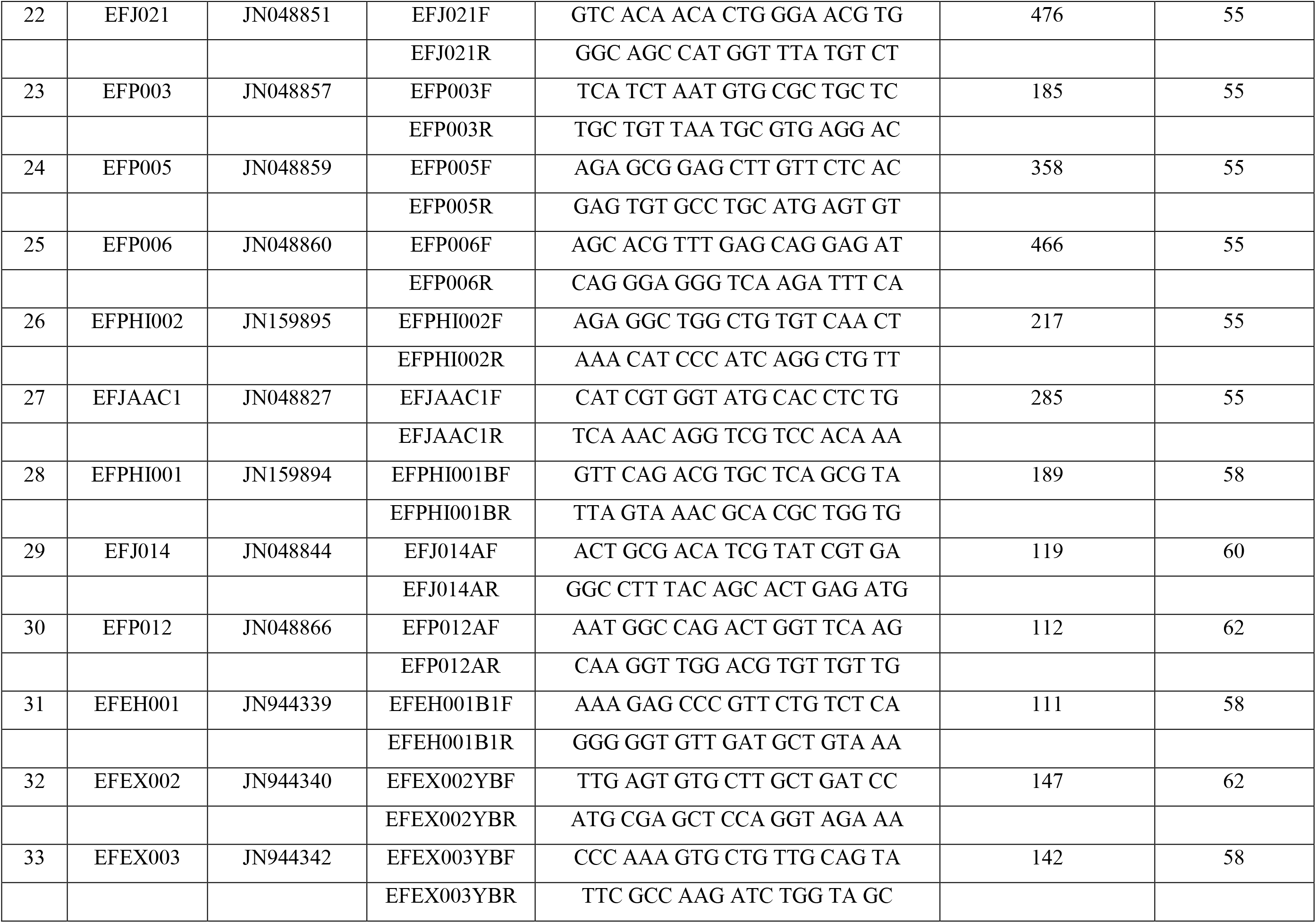
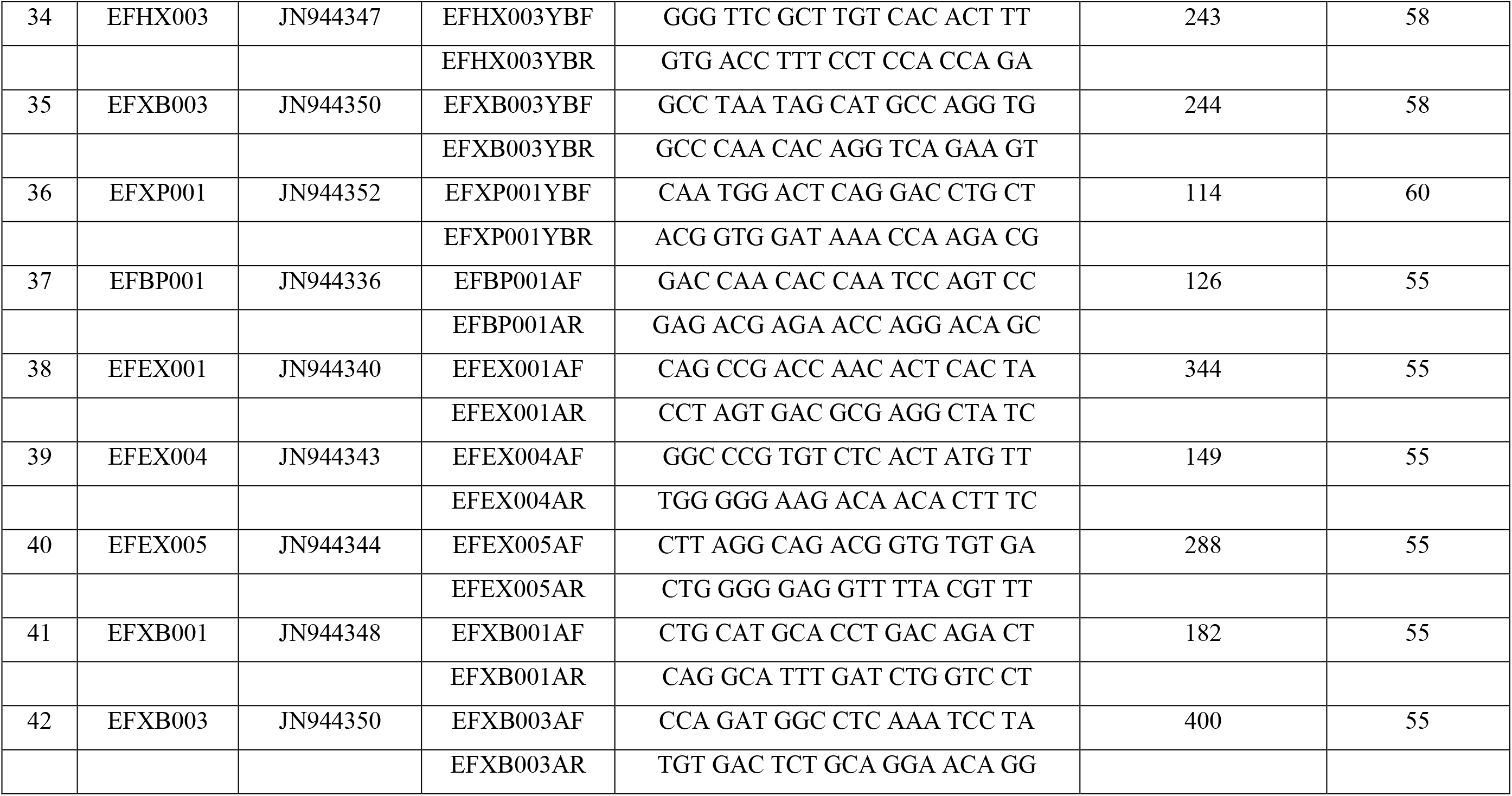
Molecular markers developed for genotyping of *E. fuscoguttatus* indicating the genomic locus, GenBank accession number, primer designation, sequences of forward and reverse primers, expected product size and annealing temperatures.

**Table 2.0:**
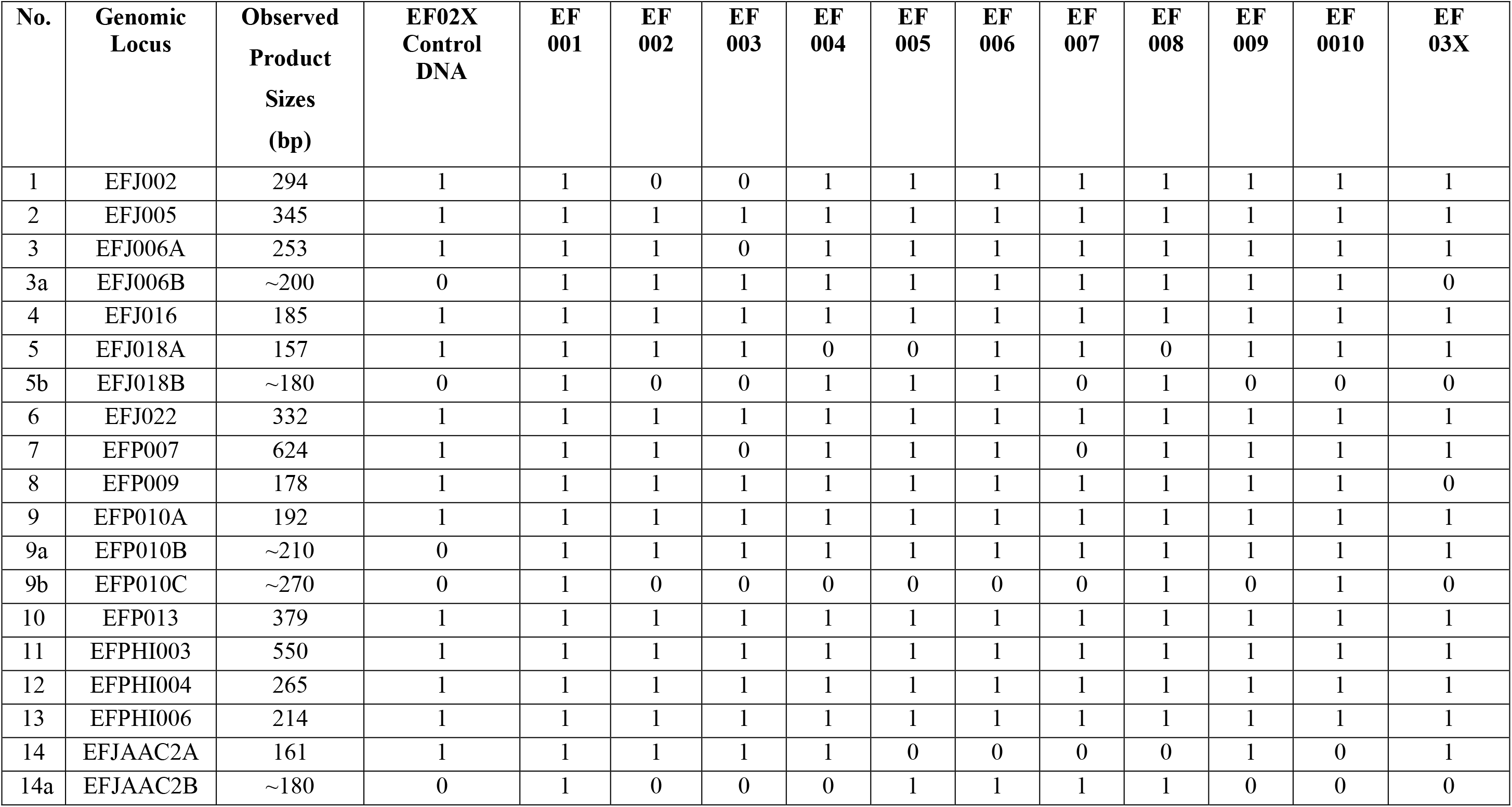
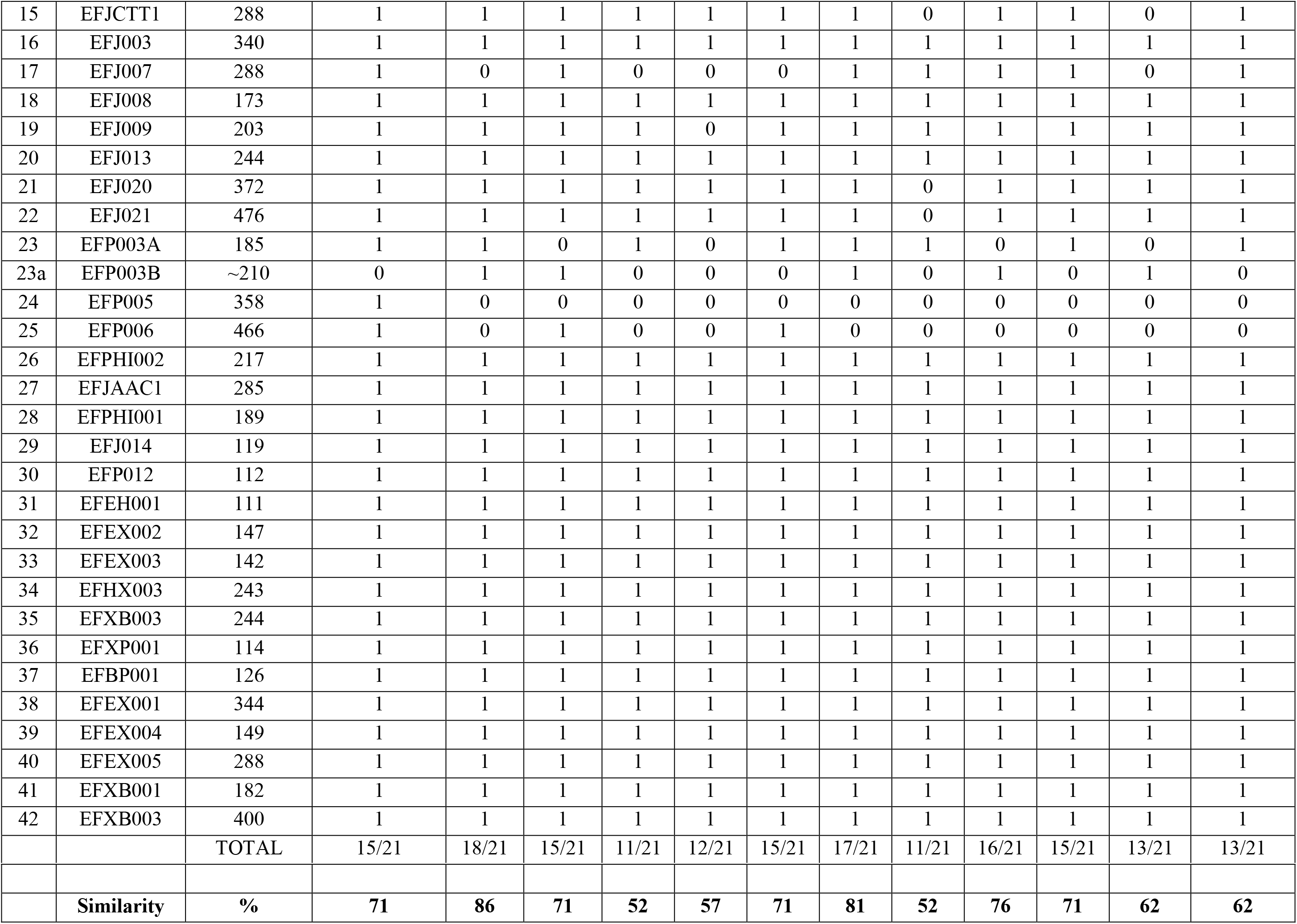
Intra-specific genetic profiling of *E. fuscoguttatus* broodstock collected from the wild. ‘1’ indicates the presence of a PCR amplicon of the expected product size, ‘0’ indicate the absence of a PCR amplicon. Primers EFJ006, EFJ018, EFP010, EFJAAC2 and EFP003 yielded more than one PCR amplicon and these were scored separately.

**Table 3.0:**
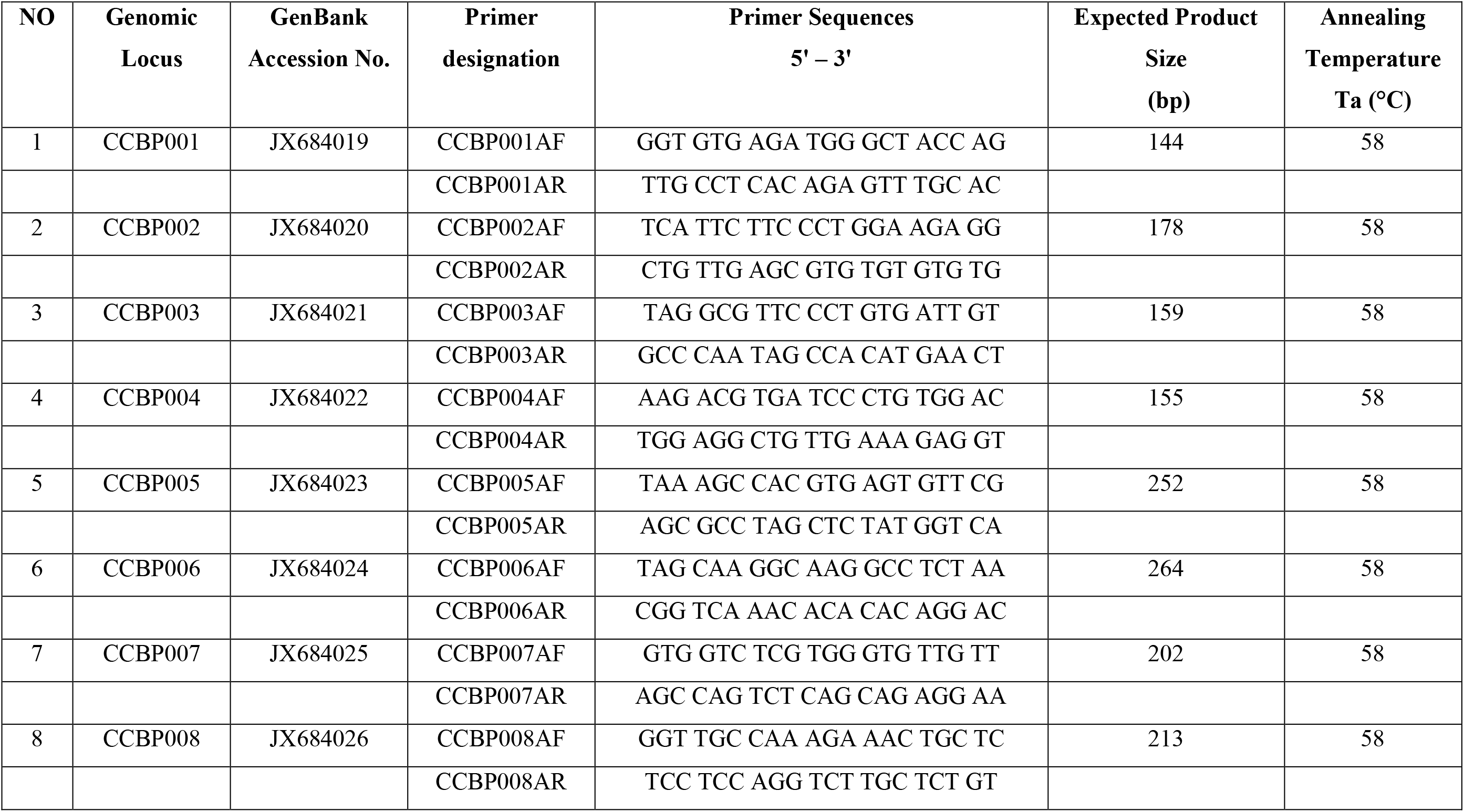
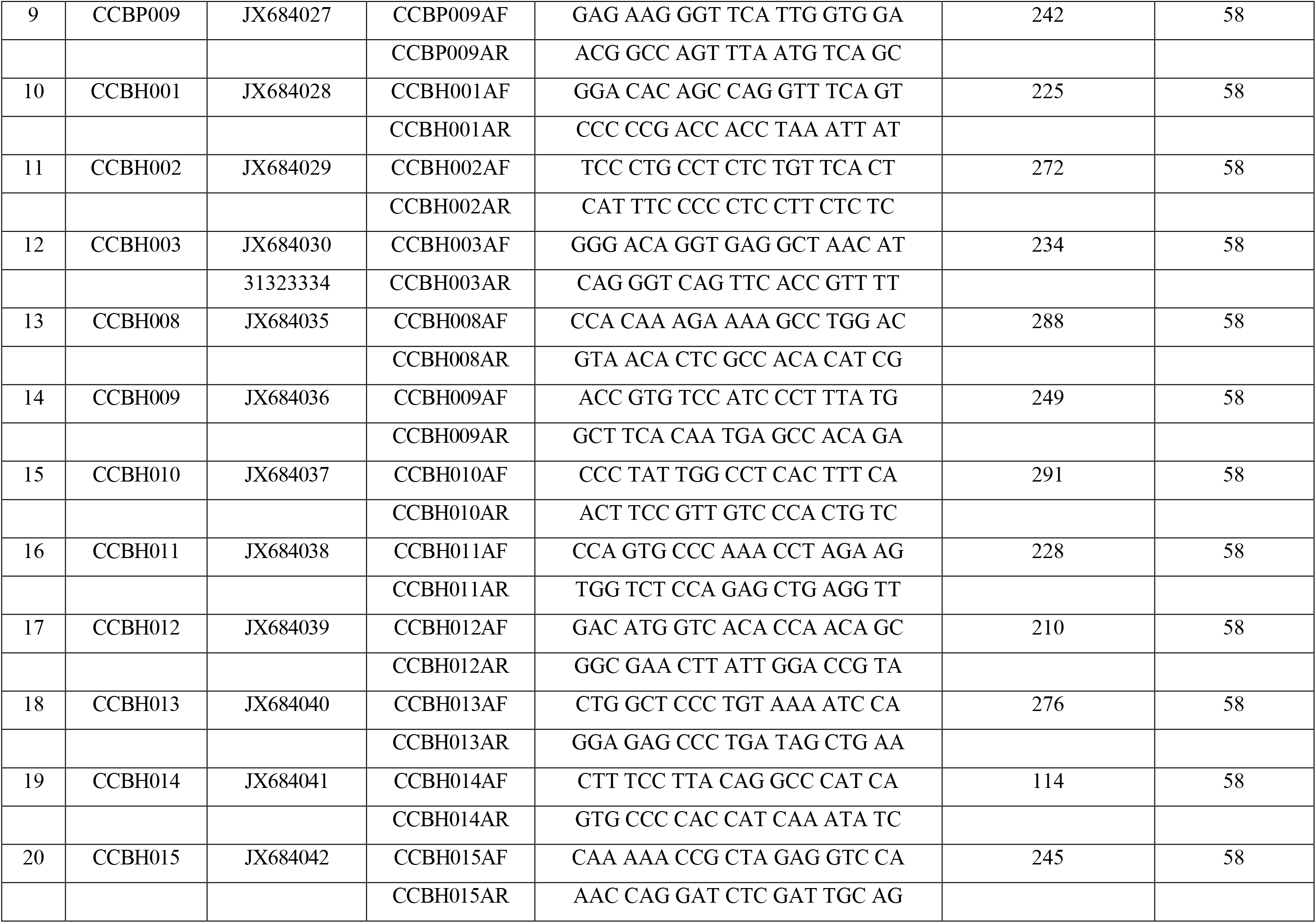
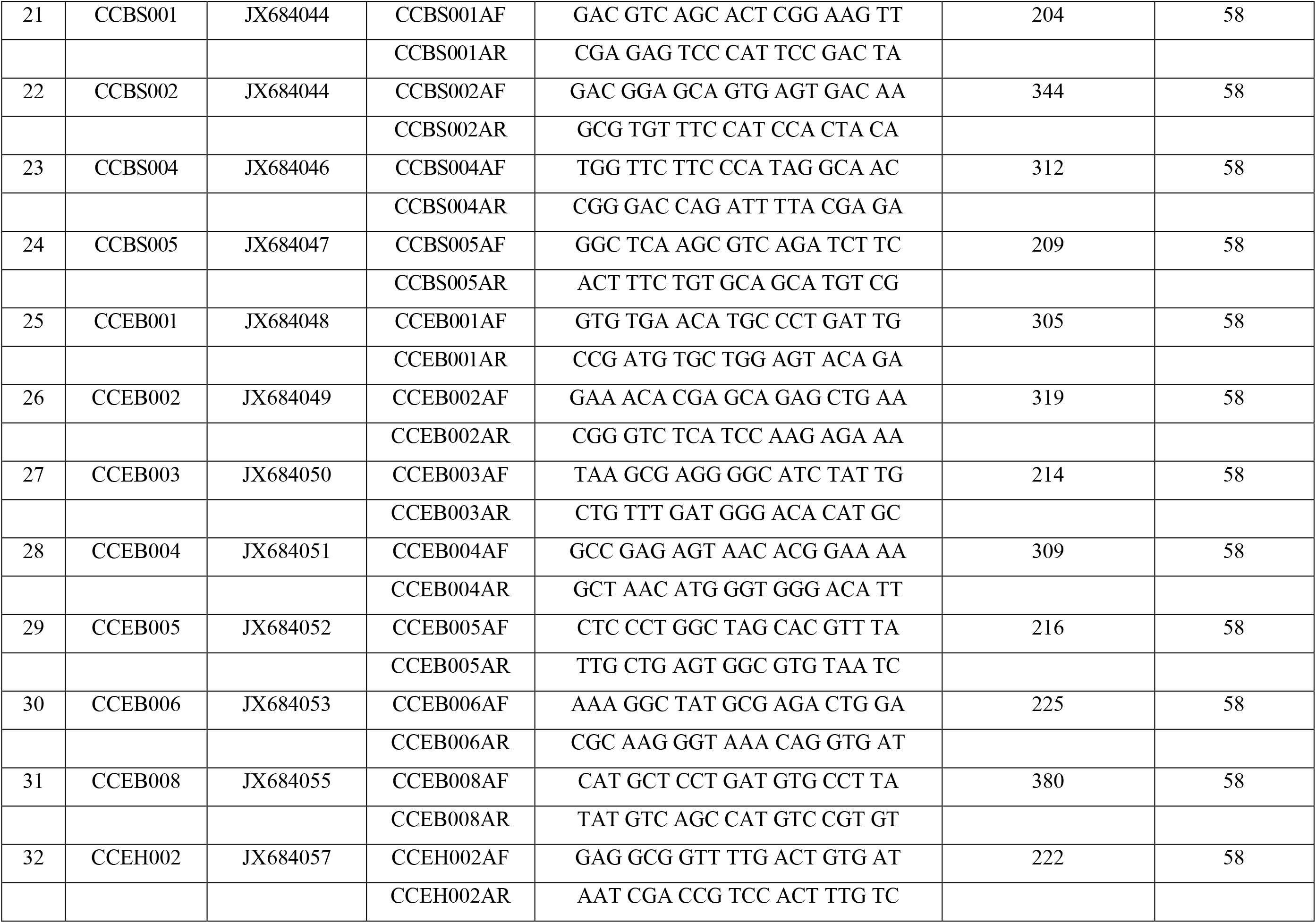
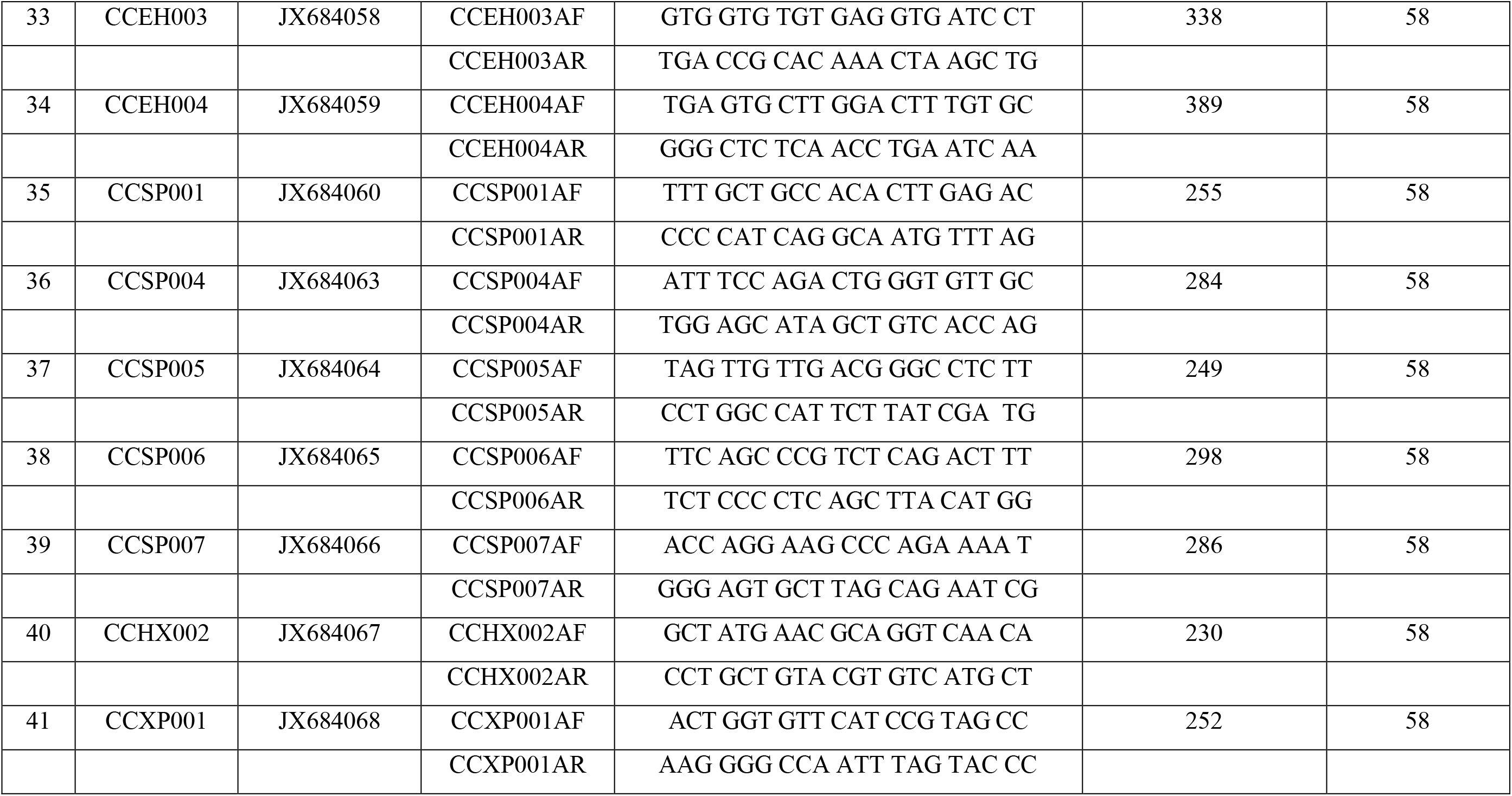
Molecular markers developed for genotyping of *E. corallicola* indicating the genomic locus, GenBank accession number, primer designation, sequences of forward and reverse primers, expected product size and annealing temperatures.

### 2.3 Primer design

One pair of primers was designed per individual DNA sequence using the online Primer3 software [16]. The annealing temperature was set (Ta) at 60 °C and regions with potential secondary structures, high G: C ratio and runs of a single nucleotide, were excluded. PCR grade primers were synthesized (IDT Technologies, Singapore).

### 2.4 Genotyping

PCR amplification was performed in final volume of 20 μl containing 1.2 μl MgCl_2_ (1.5mM), 0.4 μl dNTPs (0.2mM each), 4 μl 1x *GoTaq* buffer (Promega), 1 U *Taq* DNA polymerase (Promega), 1μl of each primer (5μM), 2μl template DNA and nuclease free water. Amplification was performed using a thermal cycler (MJ research, PTC-200) under the following conditions: pre-denaturation at 95°C for 3 min, followed by 30 cycles of denaturation (30 sec at 95°C), annealing (40 sec at 58°C), extension (2 min at 72°C) and final extension (10 min at 72°C). PCR products were resolved by electrophoresis on a 1.5% Tris-Boric Acid EDTA agarose gel, stained with Ethidium bromide and the gel was analyzed using a gel documentation system (Alpha Innotech, San Leandro, CA).

### 2.5 Scoring of PCR amplicons and data analysis

PCR amplicons were resolved on a 1.5% Tris – Boric Acid – EDTA (TBE) agarose gel, stained in a solution of Ethidium Bromide (50 μg/ml) for 10 min following which the gels were viewed using a UV transilluminator (Alpha Innotech, USA). Bands were scored as “1” for present and “0” for absent resulting in a binary score matrix. To test the reproducibility of the amplification pattern the PCR was repeated at least twice. Only unambiguous and clear amplicons were scored and chosen for Jaccard’s similarity coefficient analysis. The UPGMA phenogram was constructed using the online software DendroUPGMA (http://genomes.urv.cat/UPGMA/) [17]. Distance and Similarity matrices were computed based on the Dice Coefficient with 100 replicates. Operational Taxonomic Units (OTUs) were determined on the basis of the Cophenetic Correlation Coefficient (CPCC) [18] and phenograms were rendered graphically using the online software PhyloWidget (http://www.phylowidget.org/) [19].

## 3.0 RESULTS

### 3.1 Single Locus DNA Markers for *E. fuscoguttatus*

Six genomic libraries yielded 341 clones of which 68 were randomly selected for sequencing. Primers were designed to amplify 42 loci (Table 1.0). 37 primers yielded single amplicons whereas the remaining 5 primers (EFJ006, EFJ018, EFP010, EFJAAC2 and EFP003) amplified more than one locus. The total number of polymorphic loci was 21 (Table 2.0). The phenogram of *E. fuscoguttatus* graphically depicts the OTUs (Figure 1.0). The individuals clustered into three major clades. Clade 1 consisted of OTUs EF001, EF006, EF008, EF005, EF004 and EF010. Clade 2 consisted of OTUs EF002, EF009, EF03X, EF02X and EF003 and Clade 3 contained one OTU EF007. Within clade 1, OTUs EF001 and EF006 were grouped together with a similarity of 96%. Both of these OTUs were connected to 4 nodes containing 4 different OTUs namely; EF008, EF005, EF004 and EF010. OTU EF006 indicating highly a higher degree of similarity to two inter-joining nodes containing OTUs EF008 and EF005 with a similarity of 96% and 95% respectively. OTU EF005 was highly similar to OTU EF004 with 93% similarity. Within clade 2, OTUs EF009 and EF03X clustered with a similarity of 97%. Both OTUs were connected with three other nodes. OTU EF009 was 95% similar to OTUs EF02X, EF002 and EF003. Clade 3 was represented by only one individual OTU EGF007.

**Fig. 1.0:**
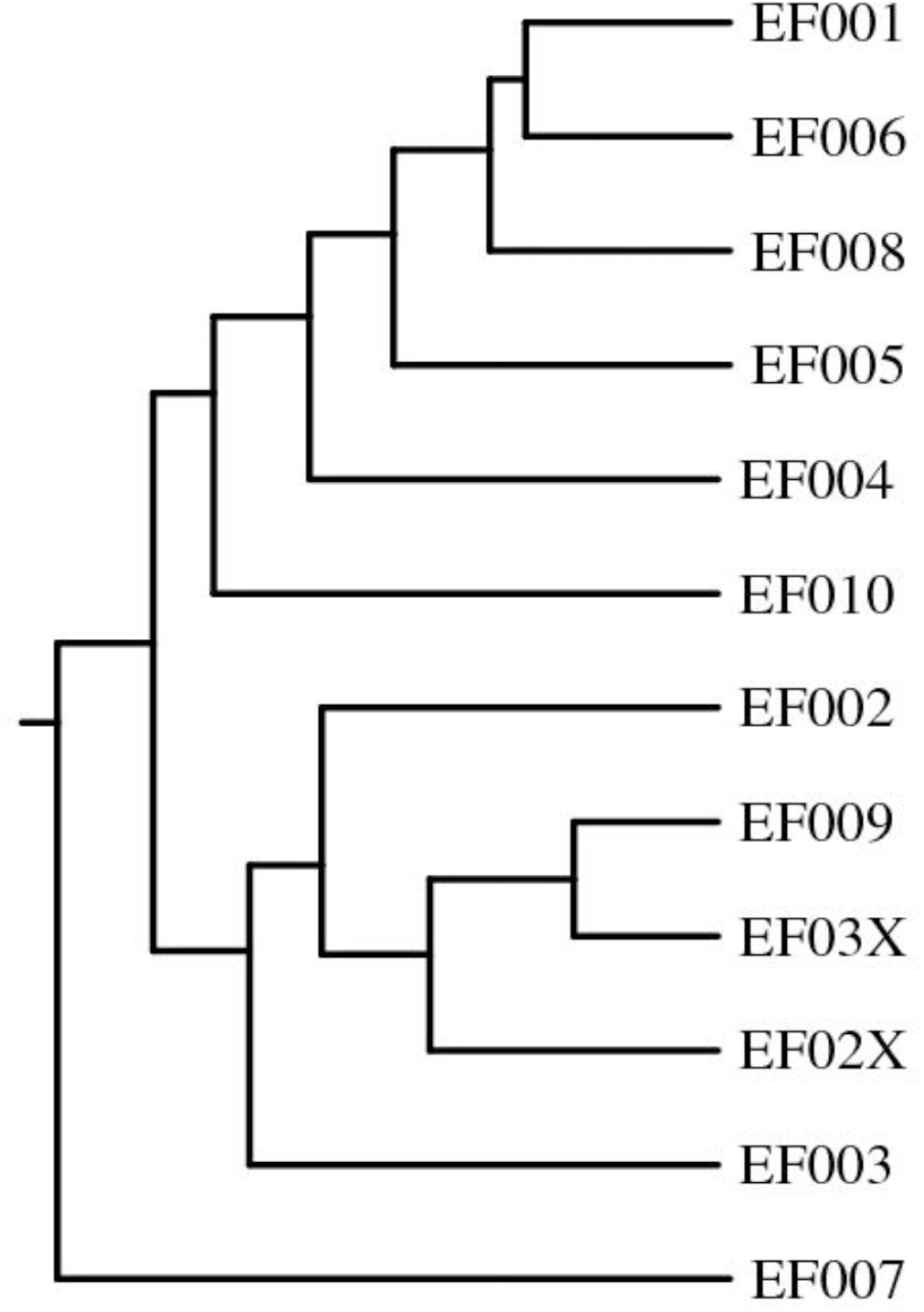
Phenogram of 12 *E. fuscoguttatus* OTU’s resulting from the UPGMA cluster analysis of the OTU x OTU correlation matrix. Cluster 1 comprises EF001, EF006, EF008, EF005, EF004 and EF010. Cluster 2 consists of EF002, EF009, EF003X, EF002X and EF003. Cluster 3 is composed on only one individual EF007.

### 3.2 Single Locus DNA Markers for *E. corallicola*

Six genomic libraries yielded 220 clones of which 52 were randomly selected for sequencing. Primers were designed to amplify 41 loci (Table 3.0) which yielded single amplicons (Table 4.0). The phenogram of *E. corallicola* graphically depicts the OTUs (Figure 2.0). The phenogram depicts three major clades. Clade 1 was composed of OTUs CC001, CC009, CC006 and CC008. Clade 2 comprised OTUs CC003, CC004, CC002, CC010, CC007 and CC005. In clade 1, OTUs CC001 and CC009 exhibited 81% similarity. Both OTUs were linked to another set of 2 nodes containing 2 different OTUs namely; CC006 and CC008.

**Figure 2.0:**
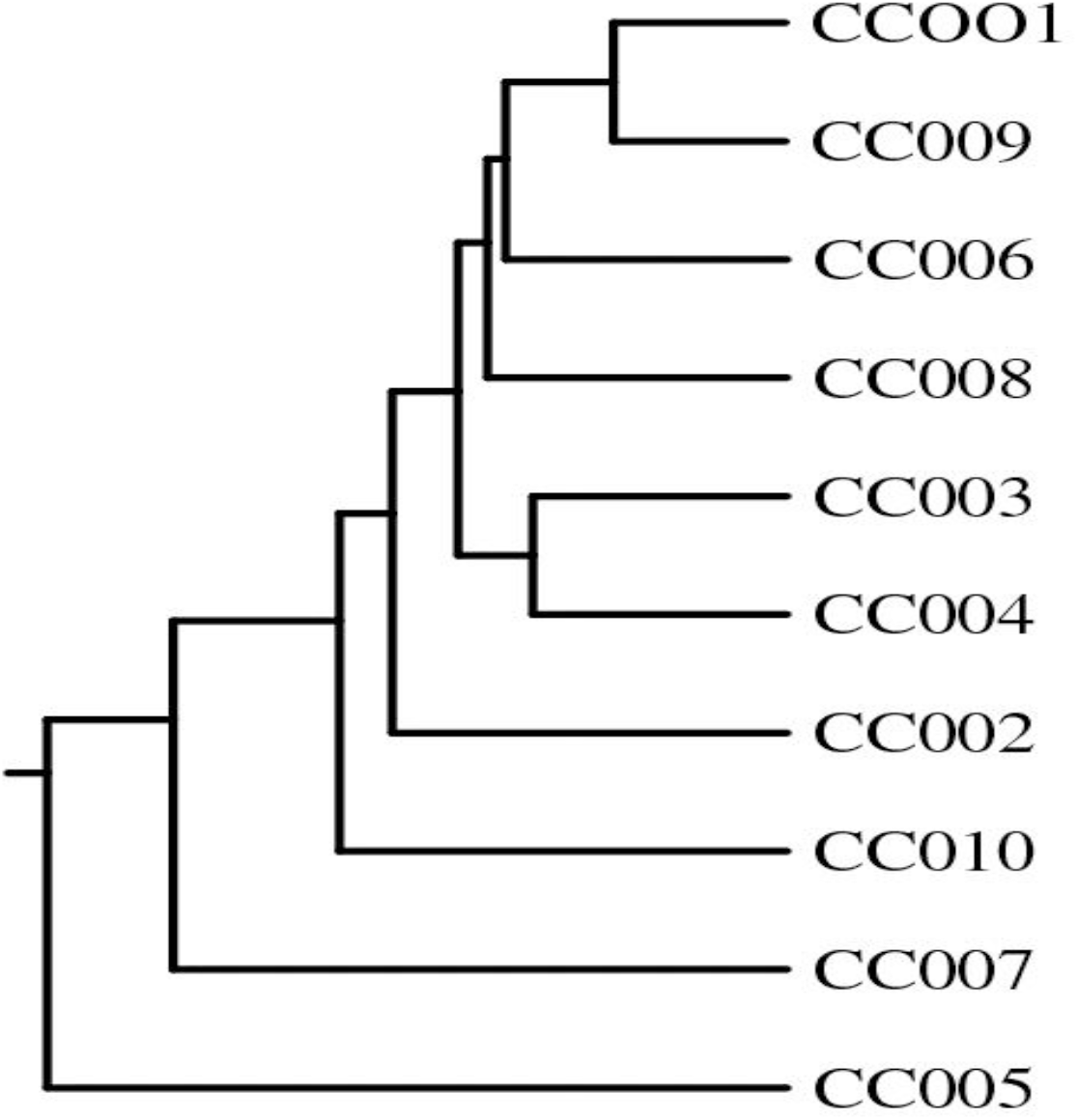
Phenogram of 10 *E. corallicola* OTU’s resulting from the UPGMA cluster analysis of the OTU x OTU correlation matrix. The high degree of intraspecific genetic diversity is evident from the distribution of the nodes.

**Table 4.0:** Intra-specific genetic profiling of *E. corallicola* broodstock collected from the wild. ‘1’ indicates the presence of a PCR amplicon of the expected product size, ‘0’ indicate the absence of a PCR amplicon.

| NO | Genomic Locus | Observed Product Size (bp) | CC001 Control | CC 002 | CC 003 | CC 004 | CC 005 | CC 006 | CC 007 | CC 008 | CC 009 | CC 010 |
| --- | --- | --- | --- | --- | --- | --- | --- | --- | --- | --- | --- | --- |
| 1 | CCBP001 | 144 | 1 | 0 | 1 | 1 | 0 | 1 | 0 | 1 | 0 | 0 |
| 2 | CCBP002 | 178 | 1 | 0 | 1 | 0 | 0 | 1 | 0 | 0 | 0 | 0 |
| 3 | CCBP003 | 159 | 1 | 1 | 1 | 1 | 0 | 1 | 0 | 0 | 1 | 0 |
| 4 | CCBP004 | 155 | 1 | 0 | 1 | 1 | 0 | 1 | 0 | 0 | 1 | 0 |
| 5 | CCBP005 | 252 | 1 | 0 | 0 | 0 | 1 | 0 | 0 | 0 | 0 | 0 |
| 6 | CCBP006 | 264 | 1 | 1 | 0 | 0 | 0 | 1 | 0 | 0 | 1 | 0 |
| 7 | CCBP007 | 202 | 1 | 0 | 0 | 0 | 0 | 1 | 0 | 0 | 1 | 0 |
| 8 | CCBP008 | 213 | 1 | 0 | 1 | 0 | 0 | 0 | 0 | 0 | 1 | 0 |
| 9 | CCBP009 | 242 | 1 | 0 | 1 | 0 | 0 | 0 | 0 | 1 | 1 | 0 |
| 10 | CCBH001 | 225 | 1 | 1 | 1 | 1 | 0 | 0 | 0 | 0 | 0 | 0 |
| 11 | CCBH002 | 272 | 1 | 1 | 0 | 0 | 0 | 1 | 0 | 1 | 1 | 1 |
| 12 | CCBH003 | 234 | 1 | 0 | 1 | 0 | 0 | 0 | 0 | 1 | 1 | 0 |
| 13 | CCBH008 | 288 | 1 | 0 | 0 | 0 | 0 | 0 | 0 | 1 | 0 | 0 |
| 14 | CCBH009 | 249 | 1 | 0 | 0 | 0 | 0 | 1 | 0 | 1 | 1 | 0 |
| 15 | CCBH010 | 291 | 1 | 0 | 0 | 0 | 0 | 0 | 1 | 0 | 0 | 0 |
| 16 | CCBH011 | 228 | 1 | 0 | 1 | 0 | 0 | 1 | 1 | 1 | 1 | 1 |
| 17 | CCBH012 | 210 | 1 | 0 | 1 | 0 | 0 | 0 | 0 | 0 | 1 | 0 |
| 18 | CCBH013 | 276 | 1 | 0 | 1 | 1 | 0 | 0 | 0 | 0 | 1 | 1 |
| 19 | CCBH014 | 114 | 1 | 0 | 1 | 1 | 0 | 1 | 1 | 0 | 1 | 0 |
| 20 | CCBH015 | 245 | 1 | 0 | 0 | 0 | 0 | 0 | 0 | 0 | 1 | 0 |
| 21 | CCBS001 | 204 | 1 | 0 | 0 | 0 | 0 | 1 | 0 | 0 | 1 | 0 |
| 22 | CCBS002 | 344 | 1 | 0 | 0 | 0 | 0 | 1 | 0 | 1 | 0 | 0 |
| 23 | CCBS004 | 312 | 1 | 0 | 0 | 0 | 0 | 0 | 0 | 1 | 0 | 0 |
| 24 | CCBS005 | 209 | 1 | 0 | 0 | 0 | 0 | 0 | 0 | 1 | 1 | 0 |
| 25 | CCEB001 | 305 | 1 | 0 | 0 | 0 | 0 | 0 | 0 | 1 | 1 | 1 |
| 26 | CCEB002 | 319 | 1 | 0 | 0 | 0 | 0 | 0 | 0 | 1 | 0 | 0 |
| 27 | CCEB003 | 214 | 1 | 0 | 0 | 0 | 0 | 1 | 1 | 1 | 1 | 1 |
| 28 | CCEB004 | 309 | 1 | 0 | 1 | 0 | 0 | 1 | 1 | 1 | 1 | 0 |
| 29 | CCEB005 | 216 | 1 | 1 | 1 | 1 | 1 | 1 | 1 | 0 | 1 | 0 |
| 30 | CCEB006 | 225 | 1 | 0 | 0 | 0 | 0 | 0 | 0 | 1 | 1 | 0 |
| 31 | CCEB008 | 380 | 1 | 1 | 1 | 1 | 0 | 1 | 0 | 1 | 1 | 0 |
| 32 | CCEH002 | 222 | 1 | 1 | 1 | 1 | 1 | 1 | 0 | 1 | 1 | 0 |
| 33 | CCEH003 | 338 | 1 | 1 | 1 | 0 | 0 | 0 | 0 | 1 | 1 | 1 |
| 34 | CCEH004 | 389 | 1 | 1 | 1 | 0 | 0 | 0 | 0 | 0 | 0 | 1 |
| 35 | CCSP001 | 255 | 1 | 1 | 1 | 1 | 1 | 1 | 1 | 0 | 1 | 1 |
| 36 | CCSP004 | 284 | 1 | 1 | 1 | 1 | 0 | 1 | 0 | 0 | 1 | 1 |
| 37 | CCSP005 | 249 | 1 | 1 | 1 | 1 | 0 | 1 | 1 | 0 | 1 | 1 |
| 38 | CCSP006 | 298 | 1 | 1 | 1 | 1 | 0 | 1 | 0 | 0 | 0 | 1 |
| 39 | CCSP007 | 286 | 1 | 0 | 0 | 0 | 0 | 0 | 0 | 0 | 1 | 0 |
| 40 | CCHX002 | 230 | 1 | 0 | 0 | 0 | 0 | 1 | 0 | 0 | 0 | 0 |
| 41 | CCHXP001 | 252 | 1 | 0 | 1 | 0 | 0 | 1 | 0 | 0 | 0 | 0 |
|  |  | TOTAL | 41/41 | 13/41 | 23/41 | 13/41 | 3/41 | 23/41 | 8/41 | 18/41 | 28/41 | 11/41 |
|  | Similarity | % | 100 | 32 | 56 | 32 | 7 | 56 | 20 | 44 | 68 | 27 |

## 4.0 DISCUSSION

### 4.1 Single Locus DNA Markers for *E. fuscoguttatus*

The tiger grouper, *E. fuscoguttatus* is highly sought after by fish breeders as it adapts well to commercial aquaculture and mariculture systems. The genetic profile indicated a high degree of similarity with 31 primers amplifying consistently across the 12 samples and 15 primers exhibiting differential profiles. This implies that the breeding population is not as genetically diverse as compared to *E. corallicola* and may require the acquisition and screening of additional recruits from the wild. Earlier reports [20] based on genotyping using microsatellites have arrived at similar conclusions regarding the abundance and diversity of wild populations of *E. fuscoguttatus*. Single locus DNA markers can be tested for Mendelian inheritance and applied for marker assisted selection of intraspecific grouper hybrids.

### 4.2 Single Locus DNA Markers for *E. corallicola*

These are the first reported genomic molecular markers for *E. corallicola*. The amplification profiles revealed a high level of intraspecific genetic diversity as the ten individuals tested exhibited unique genetic fingerprints implying that the wild population is genetically diverse. A similar study conducted using microsatellites in a closely related species *Plectropomus maculatus* led to the discovery of highly polymorphic loci which have potential applications in broodstock management [21]. Single locus DNA markers can be applied to develop linkage maps as in the case of *E. aeneus* where 222 microsatellite loci were utilized to construct a linkage map representing 24 chromosomes [22]. Advances in next generation DNA sequencing technologies have led to the development of novel markers such as SNPs as reported in *E. coioides* [23] and microsatellites with intra specific applications [24]. The development of a database of markers is essential for the development of management strategies, however markers which require extensive technical expertise and complex analysis will not be relevant to aquaculture as there is no cost benefit advantage. Under these circumstances single locus markers offer an economical alternative to SNPs.

### 4.3 Implications for broodstock management

Grouper breeders rely on wild germplasm in order to develop inbred lines. The selection of seed is done on a random basis and not on the basis of genetic diversity. There are two primary reasons for this, the first is the lack of suitable species specific markers for genetic profiling and the second is the lack of technical expertise to interpret the data derived from highly informative microsatellite DNA loci and SNPs. The objective of this study was to develop molecular markers which could be applied for diagnostic testing in a simple laboratory setup comprising a thermal cycler, gel electrophoresis system and imaging systems which are can be accessed by fish breeders. The markers which have been developed can be scored directly purely on the basis of amplification or non-amplification. Previous studies [25] that have attempted to develop molecular markers for groupers have utilized Random Amplified Polymorphic DNA (RAPD) which are difficult to reproduce due to the low degree of species specificity and high number of PCR artefacts. Furthermore, the ability of the markers to resolve intraspecific genetic polymorphism makes them ideal for population diversity studies and the construction of linkage maps for mapping quantitative traits.

## 5.0 Conclusion

This study demonstrated that species specific single locus DNA markers can be applied for the genotyping of groupers recruited from the wild for the purpose of breeding. We developed 42 and 41 novel species specific DNA markers for *E. fuscoguttatus* and *E. corallicola* respectively and applied them to resolve intraspecific diversity. These markers will be of assistance to fish breeders during the process of broodstock selection and will facilitate the process of developing genetically diverse breeding populations.

## Acknowledgements

This study was funded by the Ministry of Agriculture, Government of Malaysia, Grant Number SCF-2010-SF1010

